# Single-cell transcriptome analysis of the zebrafish embryonic trunk

**DOI:** 10.1101/2021.05.07.443129

**Authors:** Sanjeeva Metikala, Satish Casie Chetty, Saulius Sumanas

## Abstract

During embryonic development, cells differentiate into a variety of distinct cell types and subtypes with diverse transcriptional profiles. To date, transcriptomic signatures of different cell lineages that arise during development have been only partially characterized. Here we used single-cell RNA-seq to perform transcriptomic analysis of over 20,000 cells disaggregated from the trunk region of zebrafish embryos at the 30 hpf stage. Transcriptional signatures of 27 different cell types and subtypes were identified and annotated during this analysis. This dataset will be a useful resource for many researchers in the fields of developmental and cellular biology and facilitate the understanding of molecular mechanisms that regulate cell lineage choices during development.

## Introduction

The commitment of stem cells to distinct lineages is a fundamental process that underpins embryonic development. At a molecular level, a wide array of spatiotemporally regulated signaling molecules, morphogen gradients and other factors (such as physical forces) drive changes in gene expression, which guide cells down very specific lineage trajectories. Thus, understanding the dynamics of gene expression in cell populations over time is central to mapping the paths taken by cells during differentiation. Technologies such as quantitative PCR and high throughput sequencing technologies, which have emerged over the past couple of decades, have enabled scientists to probe some of these key questions in developmental biology. While traditional ‘bulk’ RNA-seq analysis can efficiently reveal transcriptional variation between different organs or organisms (e.g., wt vs. mutant), subtle changes in gene expression levels at cellular resolution cannot be achieved using this method.

In recent years, the emergence and rapid advancement of single-cell RNA sequencing (scRNA-seq) technology in combination with advances in machine learning have provided unprecedented insight into global transcriptional dynamics across different cell types [1]. The ability to capture the transcriptional information of hundreds of thousands of cells of different identities over time makes scRNA-seq an invaluable tool for dissecting cellular heterogeneity during organogenesis. Analysis of cell fate transitions at a transcriptomic level has been made possible by scRNA-seq analysis and has led to new discoveries across many fields in biomedical science. This powerful technology also has the potential to reveal transcriptomic signatures of rare and uncharacterized cell populations in disease conditions, which could revolutionize treatment strategies [2–4]. Additionally, the development of several free analytical software packages like Seurat and Monocle, which have been created to mine and analyze scRNA-seq data, has greatly facilitated research utilizing scRNA-seq [5–8].

The zebrafish (*Danio rerio*) embryo has emerged as an excellent model for studying vertebrate development due to their rapid external development, optical transparency, and high fecundity from a single mating. Furthermore, the signaling pathways that drive developmental process in zebrafish are conserved in higher vertebrates. Recent studies have used single-cell RNA-seq to characterize the transcriptomic and cellular diversity of selected tissue types [9–11]. A single cell transcriptome atlas for zebrafish embryos which encompasses one to five days of zebrafish development has been reported recently [9]. These datasets will be undoubtedly important for further studies of cell type diversity and pathways regulating choices between different lineages. However, currently available data cover only a limited number of zebrafish embryonic stages.

Here we performed single-cell transcriptomic analysis of cells isolated from the trunk region of zebrafish embryos at 30 hpf (hours post fertilization). Many important developmental processes, including definitive hematopoiesis, angiogenesis, and organogenesis are actively taking place at this time, and yet scRNA-seq at this developmental stage has not been performed previously. We focused on the trunk region because it is expected to contain all major cell types of different germ layers, including different subtypes of vascular endothelial and hematopoietic cells as well as progenitors of internal organs. At the same time, we wanted to avoid additional complexity of cell types associated with the central nervous system and craniofacial tissues which were not the focus of this analysis. 20,279 single cells were isolated from the zebrafish trunk region, resulting in a higher number of cells per cluster compared with previous analyses. We present a description of 27 transcriptionally distinct populations of cells, the identities of which were confirmed using previously published *in situ* expression data. This dataset will add to the growing database of zebrafish single-cell transcriptome data that is being generated by multiple labs in the zebrafish community. Together with previously published data, this resource will provide valuable transcriptional information on different populations of cells which could be mined and interrogated by researchers.

## Methods

### Zebrafish embryo dissociation

The zebrafish embryo study was performed under a protocol approved by IACUC at the Cincinnati Children’s Hospital Medical Center (protocol number IACUC2019-0022). Wildtype AB embryos at 30 hpf were anesthetized in 0.002% Tricaine (Sigma) and trunks of 30 embryos were dissected using a pair forceps and immediately placed in a 1.5 ml Eppendorf tube with embryo media on ice. Trunks were then dissociated into a single-cell suspension using a cold protease tissue dissociation protocol [12].

### Single-cell cDNA library preparation and computational analysis

Single cells were captured and processed for RNA-seq using the Chromium platform (10x Genomics) at the CCHMC Gene Expression Core facility. Approximately 20,279 cells were recovered with a multiplet rate of ∼7.6%. Chromium Single Cell 3’ Reagent Kits v2 was used (10x Genomics, Pleasanton, CA). 12 cDNA amplification cycles were used to generate cDNA. Sequencing parameters at a minimum were as follows: Read1, 26 cycles; i7 Index, 8 cycles; i5 Index, 0 cycles and Read2, 98 cycles. The sequencing library was sequenced at the CCHMC DNA Sequencing core on the HiSeq 2500 sequencer (Illumina, San Diego, CA) using one flow cell of paired-end 75 bp reads, generating 240-300 million total reads.

Cell Ranger version 2.2.0 was utilized for processing and de-multiplexing raw sequencing data [13]. Raw basecall files were first converted to the fastq format, and subsequently the sequences were mapped to the Danio rerio genome (version Zv11) to generate single-cell feature counts. Downstream analysis of the gene count matrix generated by CellRanger was performed in R version 3.6.0, using Seurat version 3.1 [5,14]. The gene counts matrix was loaded into Seurat and a Seurat object was created by filtering cells which only expressed more than 200 genes and filtering genes that were expressed in at least 3 cells. Additionally, as an extra quality-control step, cells were filtered out (excluded) based on the following criteria: <400 or >2,500 unique genes expressed, or >5% of counts mapping to the mitochondrial genome. This resulted in 20,279 cells in the dataset. Reads were normalized by the “LogNormalize” function that normalizes gene expression levels for each cell by the total expression, multiplies the value by a scale factor of 10^4^ and then log-transforms the result. The top 2000 highly variable genes were calculated. Prior to dimensionality reduction, a linear transformation was performed on the normalized data. Unwanted cell-cell variation driven by mitochondrial gene expression was “regressed out’ during scaling.

Dimensionality reduction was performed on the entire dataset by applying principal component analysis (PCA) using the list of highly variable genes generated above. The top 40 principal components which explained more variability (than expected by chance) were identified based on PC heatmaps, the JackStrawPlot and PCElbowPlot. 22 cell clusters were generated (by the default Louvain algorithm) using 40 PCs and a resolution of 0.5. UMAP dimensional reduction was utilized to visualize clusters [15]. Following clustering, genes differentially expressed in each of the clusters were determined using a method of differential expression analysis based on the non-parametric Wilcoxon rank

### Data availability

The original sequence files for scRNA-seq have been deposited to NCBI GEO database under accession number GSE152982.

## Results

To analyze the transcriptional profiles of different cell types in the zebrafish trunk region, the trunk portions of zebrafish embryos were manually dissected at 30 hpf as described in Methods. The cells were dissociated and subjected to single-cell RNA-seq analysis. Cells clustering resulted in the identification of 22 distinct cell populations (Fig 1, 2a). Subsequently, cell identities were assigned based on marker genes which were significantly enriched in each cluster (S1 Table and Fig 2). As an additional resource, we have also generated a table showing the expression level of all annotated genes in the zebrafish genome in all cell clusters (S2 Table).

**Figure 1:**
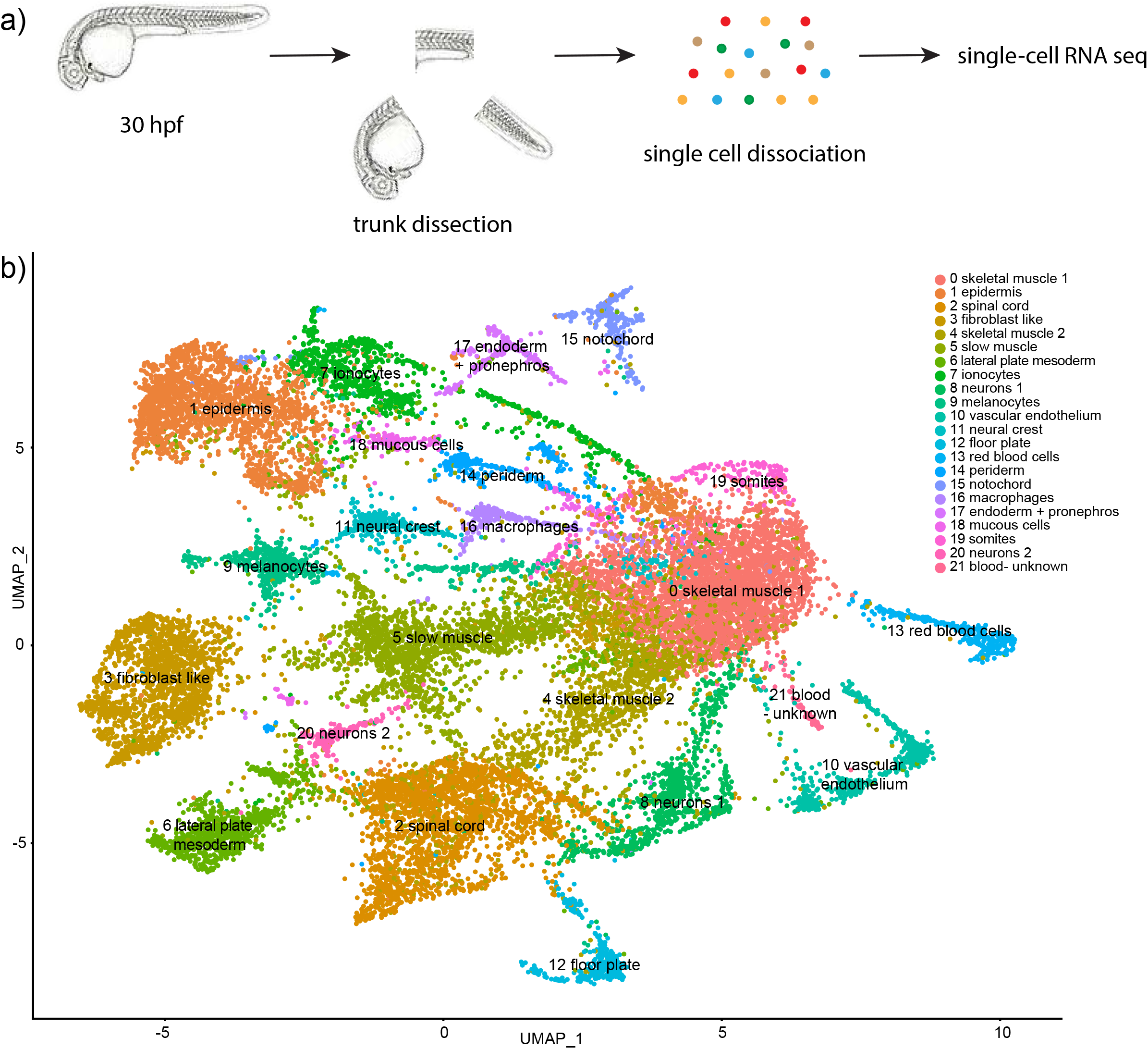
Single-cell RNA-seq analysis of trunk region in 30 hpf wild type embryos. (a) A diagram showing trunk dissection and single cell dissociation followed by single-cell RNA-seq analysis. (b) UMAP plot of 20,279 cells identified a total of 22 different cell clusters.

**Figure 2:**
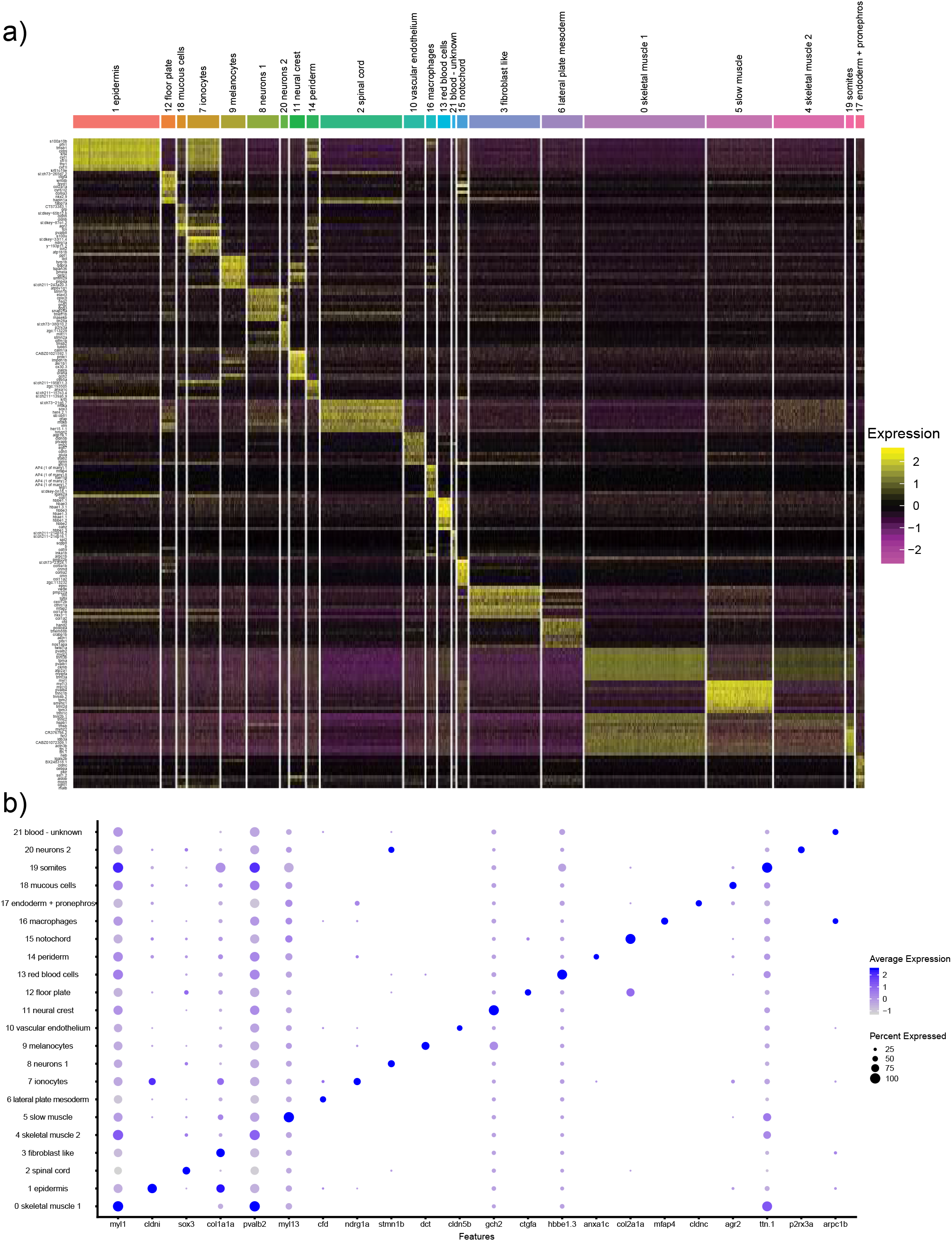
Expression of top marker genes in different cell clusters. (a) A heatmap showing expression of marker genes expression in different clusters. (b) A dot plot showing the expression of selected genes in different clusters.

### Ectodermal lineages

Nine different cell types, which included two different epidermal lineages (annotated as epidermis and periderm), ionocytes, mucous cells, melanocytes, neural crest, floor plate, spinal cord and two subsets of neurons, were identified based on differential expression analysis (S1 Table and Fig 1).

Top marker genes for epidermis (cell cluster 1), include *profilin 1* (*pfn1*), *claudin i* (*cldni*) and *keratin 4* (*krt4*) which all share expression in the zebrafish epidermis [16–18] (Fig 3a and S1 Table). Top marker genes for periderm (cell cluster 14) include *annexin A1c* (*anxa1c*), *krt4* and *krt5*, all which have reported expression patterns in the periderm or epidermis [16,17] (Fig 3b and S1 Table). There was a significant overlap of marker genes for both populations, such as *krt4* and *type I cytokeratin, enveloping layer, like* (*cyt1l*), expressed in both cell populations. Further research will be needed to determine cell identity or functional differences between the two epidermal subpopulations.

**Figure 3:**
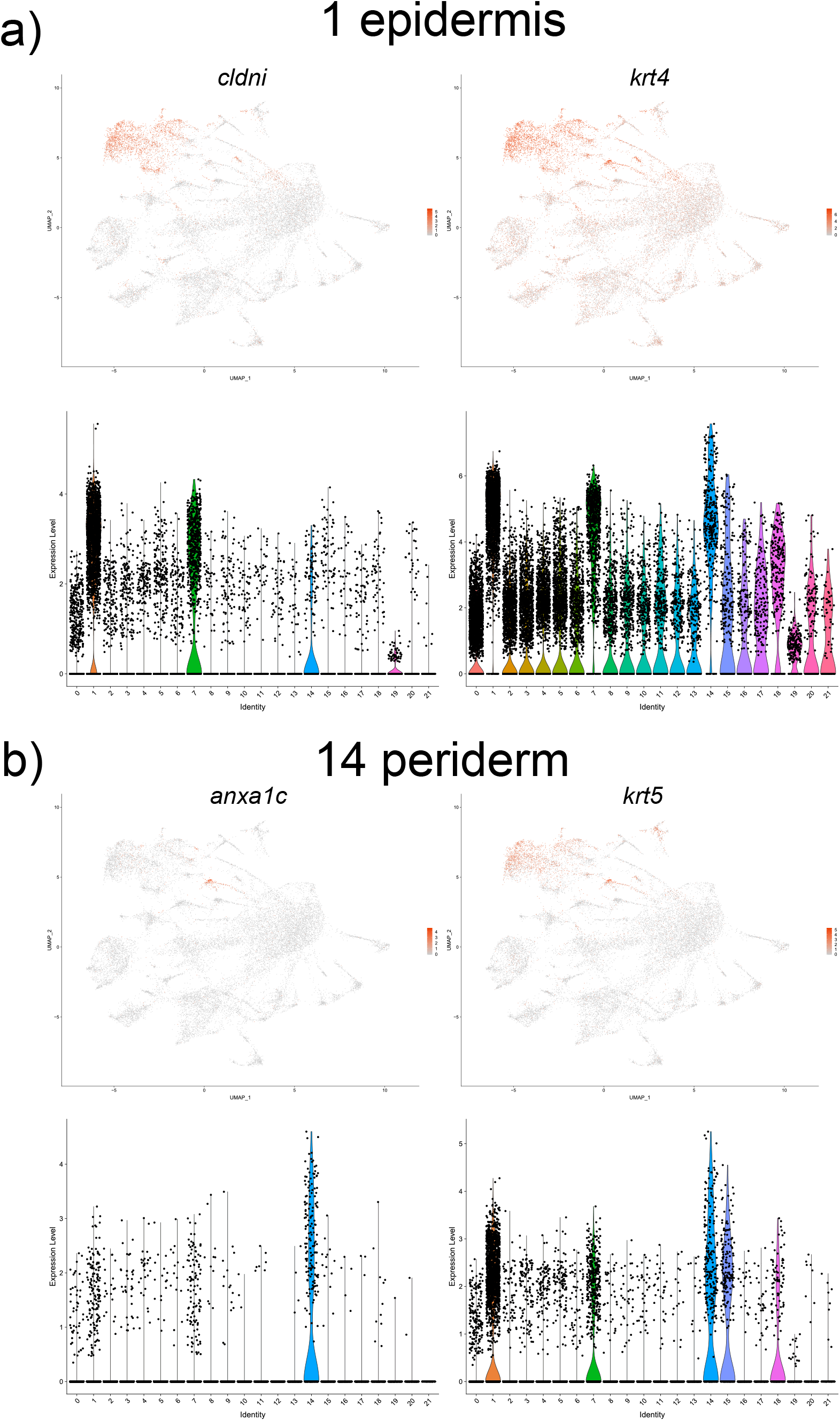
UMAP and violin plots showing the expression of: (a) *cldni* and *krt4*, top markers for epidermis (cell cluster #1); (b) *anxa1c* and *krt5*, top markers for periderm (cluster *#*14).

Top marker genes for the spinal cord (cell cluster 2) included *SRY-box transcription factor 3* (*sox3*), *hairy-related 4, tandem duplicate 2* (*her4*.*2*) and *glial fibrillary acidic protein* (*gfap*), all of which exhibit expression in the spinal cord [16] (Fig 4a and S1 Table).

**Figure 4:**
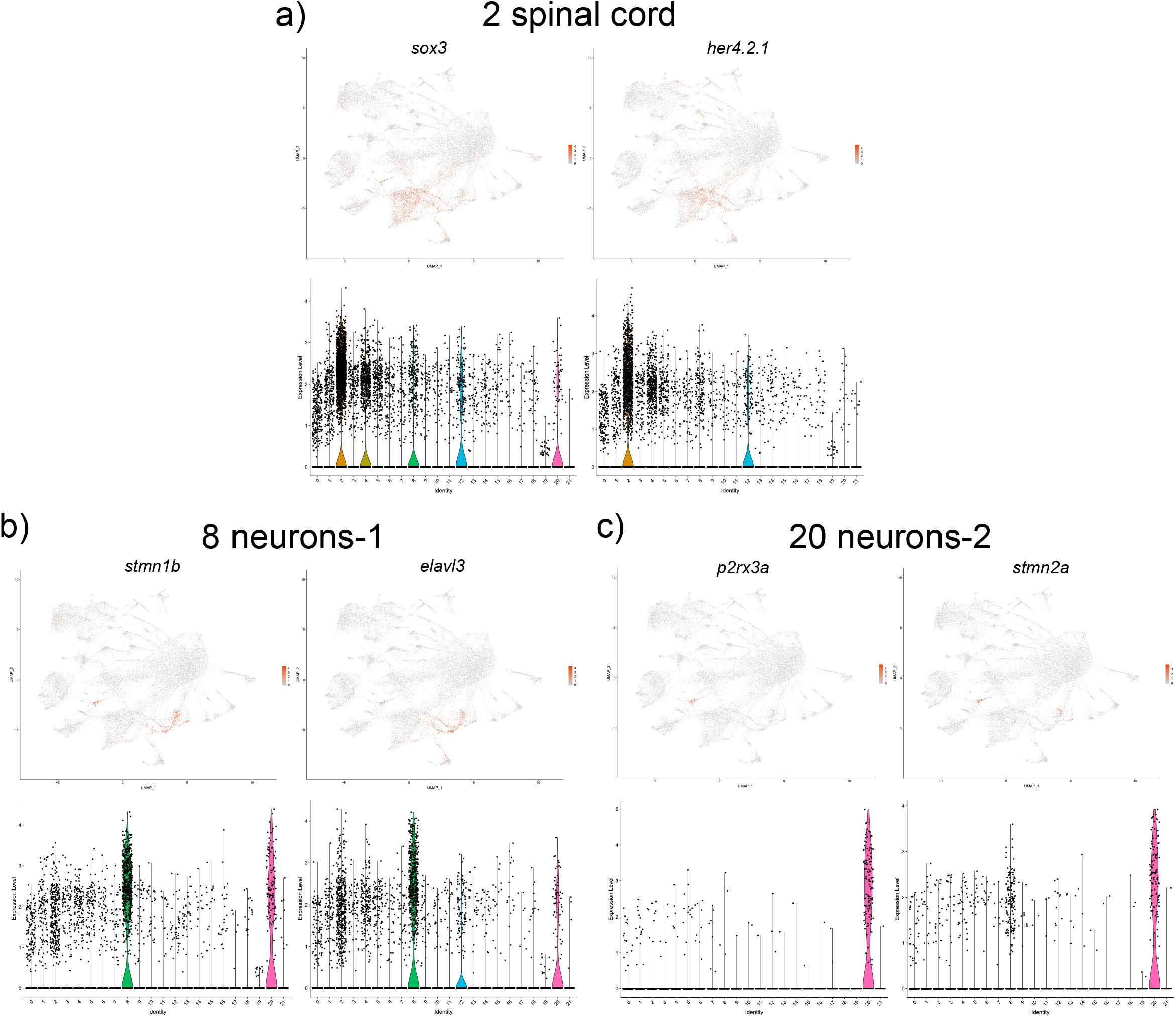
UMAP and violin plots showing the expression of selected marker genes for spinal cord (cluster #2, a), neurons-1 (cluster #8, b) and neurons-2 (cluster #20, c).

Two different cell populations with a neuronal signature were identified. Top marker genes for cell population 8 included *stathmin 1b* (*stmn1b*) and *ELAV like neuron-specific RNA binding protein 3* (*elavl3*), both of which are known neuronal markers [17,19] (Fig 4b and S1 Table). Top genes for cell population 20 included *purinergic receptor P2RX, ligand-gated ion channel, 3a* (*p2rx3a*) and *stathmin 2a* (*stmn2a)*, which both label a subset of neurons (Fig 4c and S1 Table). *p2rx3a* has been reported to label primary sensory neurons (including Rohon-Beard cells) [17,20]. While some marker genes such as *guanine nucleotide binding protein, gamma 3 (gng3)* labeled both neuronal populations, many other top marker genes were distinct between the two populations, suggesting that they represent different subtypes. It is likely that each of these populations corresponds to a specific neuronal subtype, and further investigation will be needed to characterize these subtypes.

A population of cells corresponding to melanocytes was identified (cell cluster 9); genes with high expression in this group included *dopachrome tautomerase (dct)* and *premelanosome protein a* (*pmela)* (Fig 5a and S1 Table). Both genes have been shown to be expressed in melanocytes in zebrafish [16,21]. Top marker genes for ionocytes (cell cluster 7) included *N-myc downstream regulated 1a* (*ndrg1a*) and *ATPase Na+/K+ transporting subunit beta 1b* (*atp1b1b*) [16,22] (Fig 5b and S1 Table). The top marker genes for mucus secreting cells (cell cluster 18) included *anterior gradient 2* (*agr2*) and *pvalb8*, known markers for epidermal mucus secreting cells [17,23,24] (Fig 5c, S1 Table).

**Figure 5:**
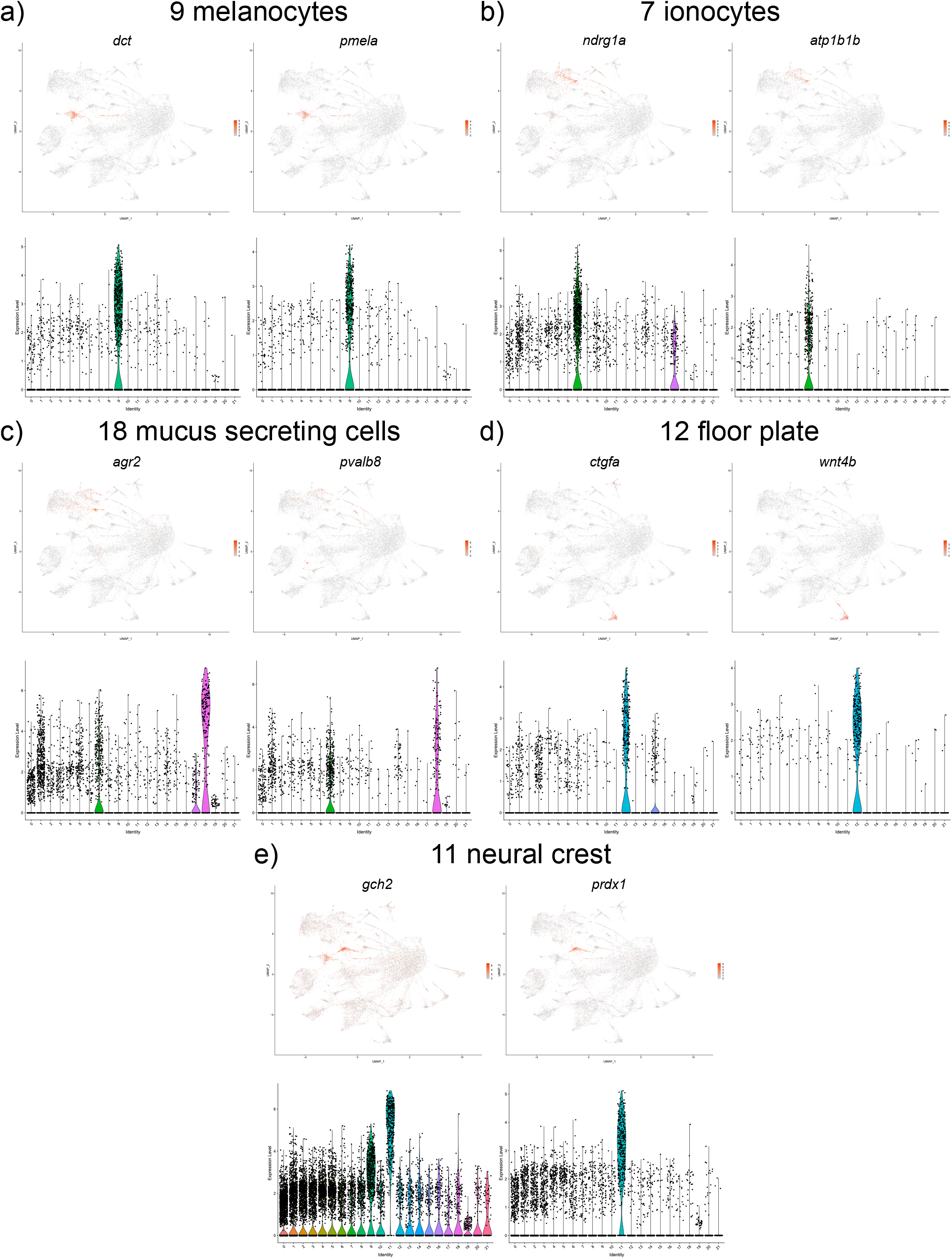
UMAP and violin plots showing the expression of selected marker genes for melanocytes (cluster #9, a), ionocytes (cluster #7, b), mucus secreting cells (cluster #18, c), floor plate (cluster #12, d) and (cluster #11, e).

The top marker genes for cell cluster 12 were *cellular communication network factor 2a (ccn2a / ctgfa)* and *wingless-type MMTV integration site family, member 4b* (*wnt4b)* which are known to be expressed in the floor plate [25,26] (Fig 5d and S1 Table). Lastly, cell cluster 11 corresponded to neural crest cells, based on the expression of its marker genes GTP *cyclohydrolase 2 (gch2)* and *peroxiredoxin 1 (prdx1*) expression [17,27] (Fig 5e and S1 Table).

### Endodermal lineage cell populations

We identified only a single cluster 17 that corresponds to the endodermal lineages. The top genes in this cell population included *claudin c (cldnc)* and *aldolase-b* (*aldob*) (S1 Table and Fig 6c). *cldnc* is a tight junction protein and *aldob* is a glycolytic enzyme involved in energy production, both of which are expressed in the digestive tract, and liver [17]. Some of the marker genes (including *cldnc* and *aldob*) also have been reported to show expression in the pronephric ducts [17],therefore we cannot exclude a possibility that pronephric cells are also included in this cluster.

**Figure 6:**
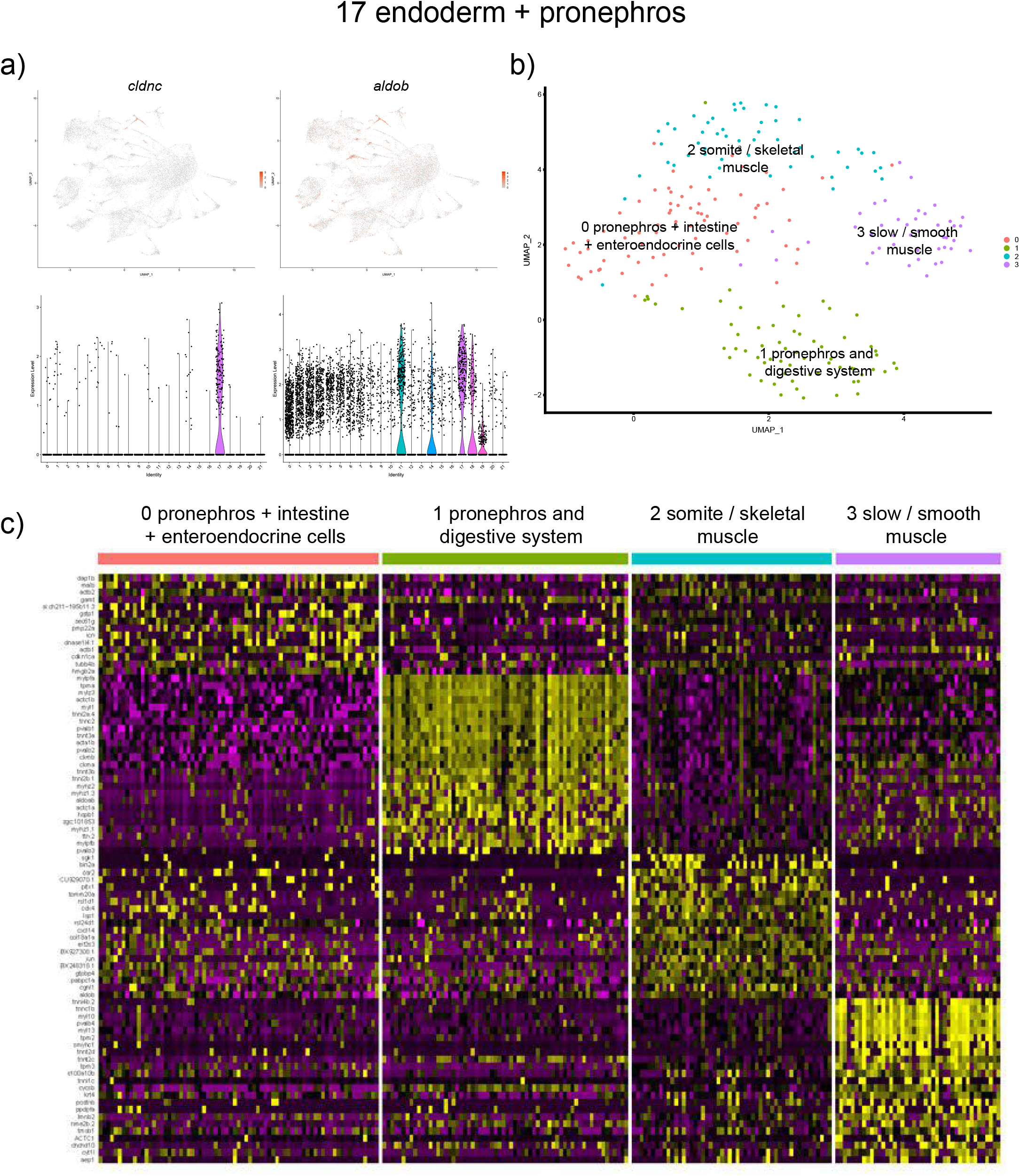
Analysis of the endodermal and pronephros cluster #17. (a) UMAP and violin plots showing the expression of top marker genes, *cldnc* and *aldob*. (b) UMAP plot showing further subclustering of this group that resulted in 4 cell subclusters. (c) A heatmap showing expression of marker genes in the subclusters.

We hypothesized that this cluster may include cells of different identities such as pronephric, liver and intestinal progenitors, thus we attempted to subcluster this cell population. Four subclusters were identified (Fig 6a,b and S3 Table); two of them included top marker genes enriched in the pronephros, intestine and / or liver. Thus, subcluster 0 included *dap1b, atp1b1a* and *selenow1* (all specific to the pronephros [16,28]) and *gstp1, agr2, gamt* (known expression in the intestine and enteroendocrine cells [29–31]). Subcluster 2 displayed top marker genes *vdrb, osr2, sgk1, pabpc1a, col18a1a* with known expression in the pronephros and digestive system [17,32–35]. Because of the significant overlap in marker expression, it was not possible to unambiguously identify these cell populations. The two remaining cell populations corresponded to somite / skeletal muscle (marker genes *mylz3, myl1*) and slow / smooth muscle identities (marker genes *myl13, myl10*). It is not entirely clear why cells with a muscle signature were mixed with this cell population, although this could reflect incomplete cell disaggregation during tissue processing.

## Mesodermal lineage cell populations

### Lateral plate mesoderm

13 different cell clusters of mesodermal origin were identified. Several clusters corresponded to lateral plate mesoderm-derived lineages. The top genes from cell cluster 10 included *aquaporin 1a*.*1 (aqp1a*.*1)* and *claudin 5b (cldn5b)*, both expressed in vascular endothelial cells [36] (Fig 7a).

**Figure 7:**
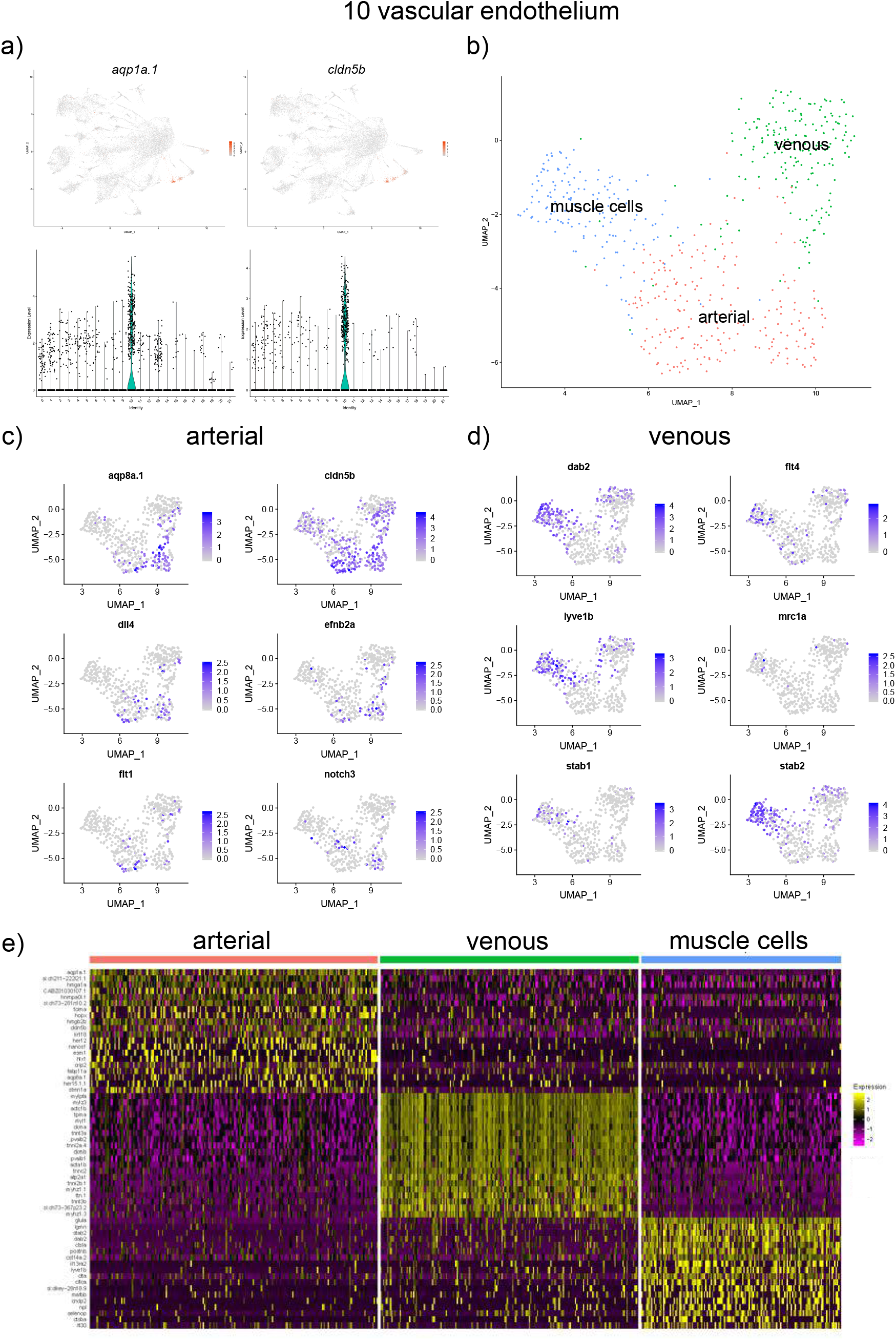
Analysis of the vascular endothelium cluster #10. (a) UMAP and violin plots showing the expression of top marker genes, *aqp1a*.*1* and *cldn5b*. (b) UMAP plot showing further subclustering of this group. 3 subclusters which show transcriptional signature of arterial, venous and muscle cells were identified. (c,d) UMAP plot showing the expression of selected arterial and venous specific markers within the vascular endothelial cluster. (e) A heatmap showing expression of marker genes in the subclusters.

Vascular endothelial cells are known to exhibit substantial diversity. Arterial, venous, lymphatic progenitors can be distinguished as early as 24 hpf or even earlier stages [37]. Because only a single vascular endothelial cluster was identified during the initial clustering, we performed further subclustering of vascular endothelial cells. Three different subpopulations were identified, which included arterial and venous cells, based on the signature of marker genes (*aqp1*.*a1, aqp8a*.*1, cldn5b, hey2* and others for arterial cells; *stab2, dab2, lyve1b* for venous cells) (Fig 7b-e). Multiple other marker genes were identified in each population, some of which are likely to be novel arterial and venous marker genes (S4 Table). The third population was enriched in marker genes specific for muscle cells, including *pvalb2, mylz3, myl1* and others. Intriguingly, the expression of vascular endothelial-specific genes such as *cdh5* was also observed within this population. Previous studies have argued that somites contribute to vascular endothelial cells [38–41]. It is possible that this cell population corresponds to these transitional somite-derived endothelial cells. Alternatively, we cannot exclude the possibility that some perivascular cells were tightly attached and failed to separate during cell dissociation, resulting in doublets of mixed identity.

We identified three different blood cell clusters from our analysis. Cell cluster 13 corresponds to red blood cells (RBCs), based on the marker gene hemoglobin, beta embryonic 1.3 (*hbbe1*.*3)* and hemoglobin alpha embryonic-3 (*hbae3)* expression [18,42– 44] (S1 Table and Fig 8a). The top marker genes for cell cluster 16 included *microfibril associated protein 4 (mfap4)* and *lysozyme g-like 1 (lygl1)*, both known to label macrophages [17,45] (S1 Table and Fig 8b). Lastly, cell cluster 21 appears to correspond to a distinct hematopoietic population. The expression of top marker genes including *cd59* and *actin related protein 2/3 complex, subunit 1B (arpc1b)* (S1 Table and Fig 8c) has been reported in blood cells [46,47]. Further studies will be needed to confirm the identity of this population.

**Figure 8:**
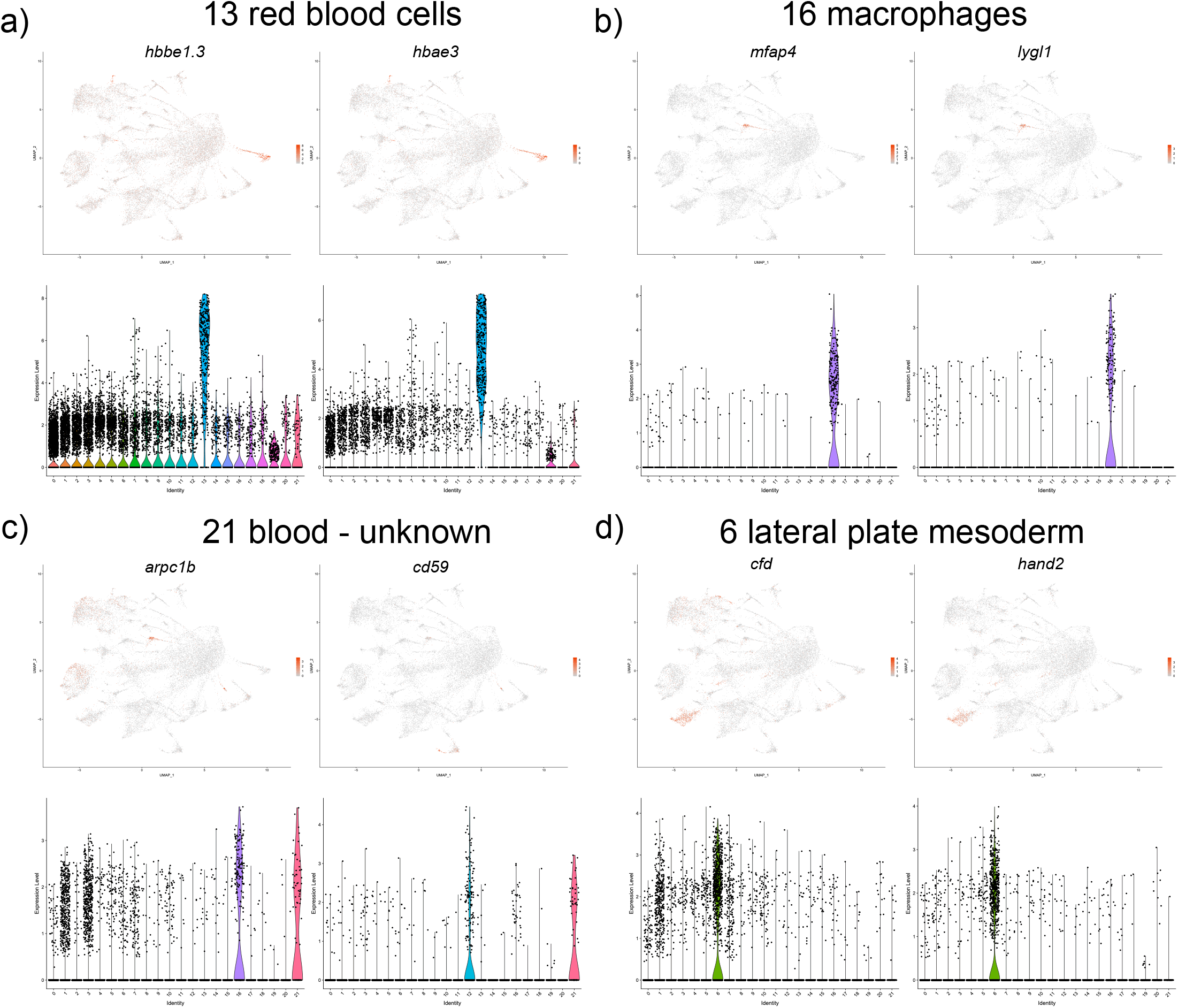
UMAP and violin plots showing the expression of selected marker genes for red blood cells, (cluster #13, a), macrophages (cluster #16, b), blood cells of unknown identity (cluster #21, c) and lateral plate mesoderm (cluster #6, d).

Cell cluster 6 corresponds to a poorly understood cell population that is likely derived from the lateral plate mesoderm. Its top marker genes include *complement factor D (cfd), heart and neural crest derivatives expressed 2 (hand2)* (S1 Table and Fig 8d), which are known to have bilateral expression along the yolk sac extension at this stage (30 hpf) [17,48]. At earlier somitogenesis stages, *hand2* has prominent expression in the LPM [48]. Some additional top markers genes in this population such as *twist1a* also share LPM expression during somitogenesis stages [49].

### Paraxial and axial mesoderm

Two different subsets of fast skeletal muscle were identified. The top markers in cell cluster 4 included *myosin, light chain 1, alkali; skeletal, fast (myl1)* and *heat shock protein, alpha-crystallin-related, 1 (hspb1)*, both known to be expressed in skeletal muscle cells [50,51] (S1 Table and Fig 9a). Cluster 0 also had multiple genes expressed in the myotome and skeletal muscle cells including parvalbumin 2 (pvalb2) and *myosin, light polypeptide 3, skeletal muscle (mylz3)* [24,52] (S1 Table and Fig 9b). There was a significant overlap in gene expression between the two groups of muscle cells, and the marker genes *myl1, mylz3, pvalb2* were among the top ten marker genes for both cell groups. Cell cluster 5 was characterized by the expression of top marker genes *myosin, light chain 13 (myl13)* and *myosin, light chain 10 (myl10)*, known markers of slow muscle cells [52] (S1 Table and Fig 9c). Cell cluster 19 included marker genes *titin (ttn*.*1 and ttn*.*2)*, and *actinin alpha 3b (actn3b)*, known to be expressed in the somites and skeletal muscle cells [53,54] (S1 Table and Fig 9d). Cell cluster 3 included marker genes *transforming growth factor, beta-induced (tgfbi)* and collagen *col1a1b* which have known expression in fibroblast-like cells present at myotendinous junctions (S1 Table and Fig 10a). And lastly, cell cluster #15 corresponded to the notochordal cells based on the expression of top marker genes *collagen col2a1a* and *col9a1b* [16,55] (S1 Table and Fig 10b).

**Figure 9:**
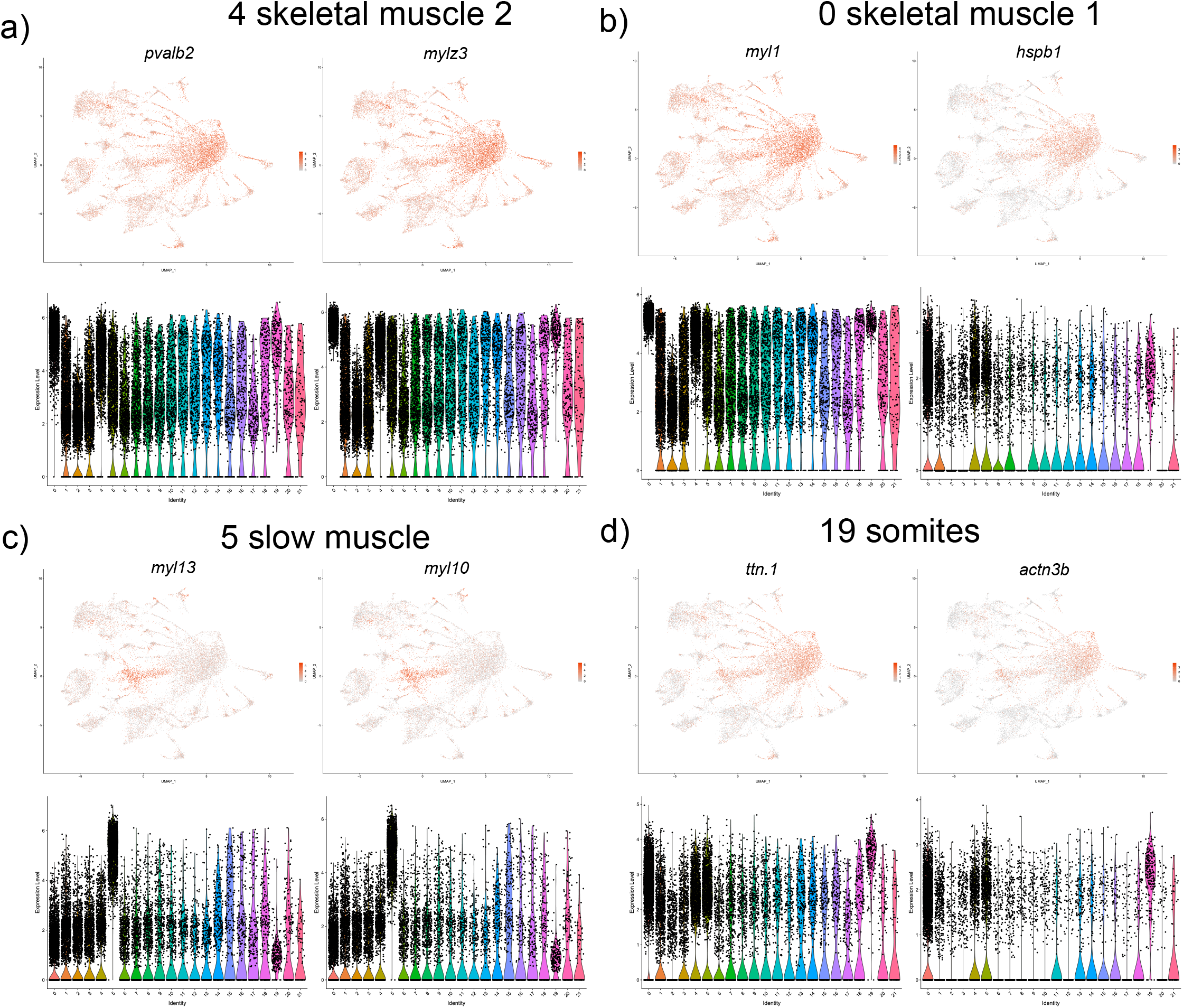
UMAP and violin plots showing the expression of selected marker genes for two skeletal muscle groups (clusters #4 and #0, a,b), slow muscle (cluster #5, c) and somites (cluster #19, d).

**Figure 10:**
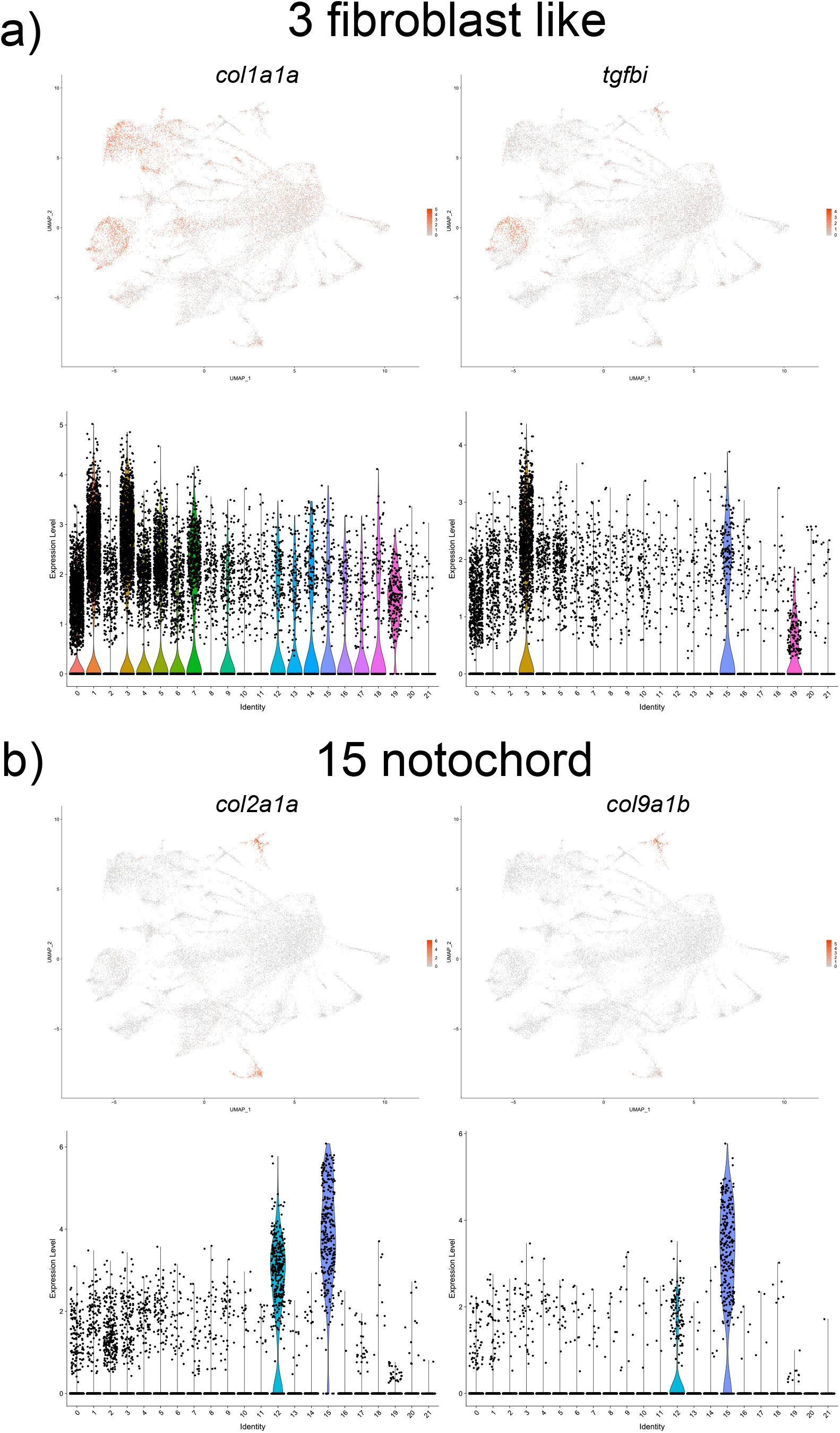
UMAP and violin plots showing the expression of selected marker genes for fibroblast like (cluster #3, a) and notochordal cells (cluster #15, b).

## Discussion

In the current study we have identified 22 distinct cell clusters two of which were subclustered resulting in total of 27 cell groups with unique transcriptional signatures. Many of the groups had unique and easily identifiable signatures (endothelial cells, melanocytes, macrophages and others). In some of the groups there was a significant overlap in marker expression. For example, skeletal muscle 1 and 2 groups (clusters 0 and 4) share expression of many top markers, including *pvalb2, mylz3, tpma* and others. At the same time, many genes also show differential expression between the two groups. Further studies will be needed to validate biological differences between these populations. And lastly, some cell clusters had poorly characterized identities, including lateral mesoderm and blood – unknown (clusters 6 and 21). Many of the top markers of cluster 21 have no or poorly characterized expression in zebrafish. Some of the marker genes, including *lmo2, myb* and *fli1a* are known to be expressed in hematopoietic stem cells [56]. *arpc1b* function in mouse has been implicated in T-cell and thrombocyte development [46]. Further in-depth analysis is required to analyze these cell populations.

A recent study has reported a single-cell transcriptome atlas for zebrafish, based on scRNA-seq analysis of whole zebrafish embryos at 1, 2 and 5 dpf stages [9]. 220 different clusters were described based on the analysis of 44,102 cells in this study. Our analysis was limited to the trunk region at 30 hpf stage which is a different stage than previously analyzed, therefore it is difficult to compare these datasets directly. Although Farnsworth *et al* have reported significantly more cell clusters, it is unclear yet if all of them represent truly distinct cell types. Also, our analysis was limited to a single stage at the trunk region, therefore fewer cell groups are expected. Some of the clusters identified in our study such as arterial and venous specific transcriptomes, or the unknown blood group (cluster 21) have not been reported in the previous study.

In summary, our results provide a unique resource for cell lineages located in the trunk region of a developing zebrafish embryo and will complement transcriptomic datasets generated by other groups. This information will be essential in deciphering the signaling pathways and transcriptional programs that regulate the establishment and differentiation of a variety of cell types during vertebrate development.

## Acknowledgments

We thank Andrew Potter and Steve Potter for single cell dissociation protocol and assistance, Shawn Smith and Kelly Rangel (Gene expression core – CCHMC) for assistance in processing samples for scRNAseq, Praneet Chaturvedi for his assistance with scRNA-seq analysis.

## Author contributions

S.M. performed zebrafish embryo dissociation and wrote the manuscript, S.C.C. performed scRNA-seq data computational analysis and wrote the manuscript, S.S. conceived and supervised the project and edited the manuscript.

## Competing statements

Authors declare no competing interests.

## Supporting information

**S1 Table. Differential expression of marker genes in different cell clusters. S2 Table. Average gene expression in different cell clusters**.

**S3 Table. Differential expression of marker genes in endoderm + pronephros subcluster - #0 pronephros + intestine + enteroendocrine cells**.

**S4 Table. Differential expression of marker genes in endothelial cell subcluster - #0 arterial**.

